# Minimal integrating shuttle vectors for *Saccharomyces cerevisiae* depleted of restriction sites outside the polylinker region

**DOI:** 10.1101/2024.11.05.622133

**Authors:** Lorenzo Scutteri, Patrick Barth, Sahand Jamal Rahi

## Abstract

Many plasmids harbor unnecessary elements that complicate or hinder cloning tasks such as inserting one gene into another for protein domain grafting. In particular, restriction sites may be present in the backbone outside the polylinker region (multiple cloning site; MCS) and thus unavailable for use, and the overall length of a plasmid correlates with poorer ligation efficiency. To address these concerns, there has been a growing interest in minimal plasmids. Here, we describe the design and validation of a collection of six minimal integrating shuttle vectors for genetic manipulation in *Saccharomyces cerevisiae*. We constructed the plasmids using *de novo* gene synthesis and consisting only of a yeast selection marker (*HIS3, TRP1, LEU2, URA3, natMX6*, or *KanMX*), a bacterial selection marker (Ampicillin resistance), an origin of replication (ORI), and the MCS flanked by M13 forward and reverse sequences. We use truncated variants of these elements where available and eliminated all other sequences typically found in plasmids. The MCS consists of ten unique restriction sites. To our knowledge, at sizes ranging from approximately 2.6 kb to 3.5 kb, these are the smallest shuttle vectors described for yeast. Further, we removed common restriction sites in the open reading frames (ORFs) and terminators, freeing up approximately 30 cut sites in each plasmid. We named our pLS series in accordance with the well-known pRS vectors, which are on average 63% larger: pLS403 (*HIS3*), pLS404 (*TRP1*), pLS405 (*LEU2*), pLS406 (*URA3*), pLS408 (*natMX6*), and pLS410 (*KanMX*). These minimal vector backbones open up new opportunities for efficient molecular biology and genetic manipulation in *Saccharomyces cerevisiae*.

## INTRODUCTION

Plasmids are important tools in molecular biology, needed for genetic engineering and recombinant gene expression in various host organisms^1–3^. Yeast shuttle vectors generally range from 4 to 10 kb in size^4–15^ and harbor superfluous elements and restriction sites outside the MCS that complicate experiments. Larger plasmids tend to be more difficult to manipulate, for example, when needing to clone multiple genes into the same backbone^16^. Certain applications, e.g., DNA break repair assays, require avoiding the same DNA sequences around engineered genomic modifications.^17^ Also, restriction sites that are present in the backbone cannot be used to modify insert DNA. For example, for grafting a protein domain into another, many different insertions sites typically need to be tested, requiring as few restriction sites as possible to be used up in the vector backbone.

To address these limitations, there has been a growing interest in developing size-reduced plasmids. This minimalistic philosophy aims to create compact, precisely engineered vectors that only contain essential elements. By reducing plasmid size and eliminating unnecessary sequences, minimal plasmids have been shown to enhance cloning efficiency, gene transfer, and genetic manipulation^18–20^. Size-reduced cloning vectors have been previously constructed through the progressive miniaturization of existing commercial plasmids, but an aggressive miniaturization strategy has not been applied to shuttle vectors for *Saccharomyces cerevisiae*^21–24^.

In the present study, we leveraged increasingly affordable *de novo* gene synthesis to create minimal shuttle vectors for *Saccharomyces cerevisiae*. The pLS plasmids only contain a yeast selection marker (*HIS3, TRP1, LEU2, URA3, natMX6*, or *KanMX*), the bacterial selection marker *AmpR* conferring Ampicillin resistance, an ORI, and a Multiple Cloning Site (MCS) flanked by M13 forward and reverse sequencing regions (Fig. 1A-F). We used truncated variants of these elements where available and avoided unnecessary DNA sequences. Furthermore, we recoded the yeast selection markers and *AmpR*, and we mutated terminators to enhance the availability of restriction sites for insert manipulation. We introduced, on average, 68 mutations per plasmid, removing most cut sites in the vector backbone while preserving the original amino acid sequences. Recoding the selectable markers recovered approximately 30 restriction sites recognized by widely used Type IIP and IIS restriction endonucleases in each plasmid.

**Figure 1.**
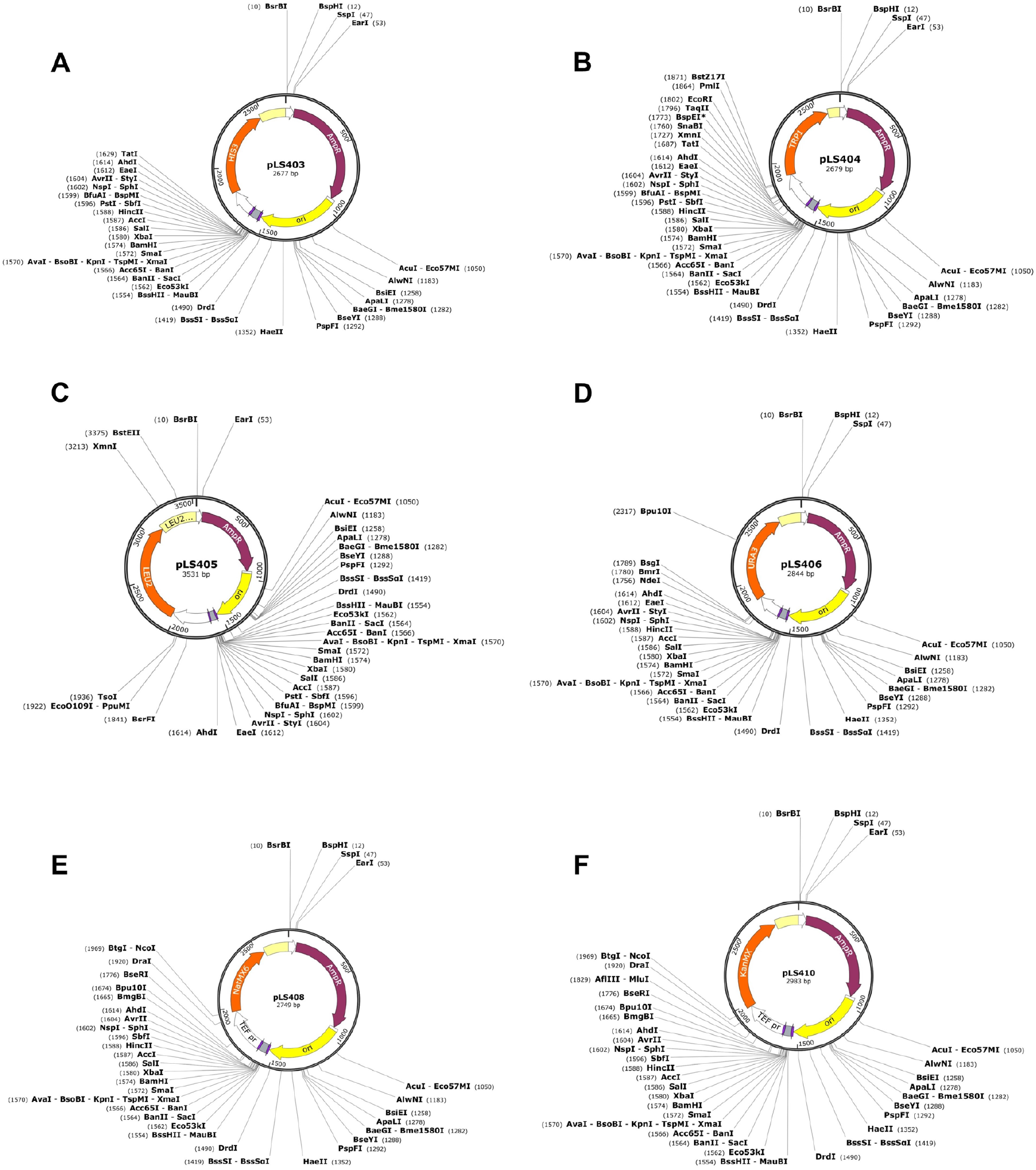
Maps of the pLS minimal shuttle vectors. For each minimal shuttle vector, the size and the unique restriction sites are indicated in the maps.

We named the pLS plasmids in accordance with the well-known pRS400 series^5^: pLS403 (*HIS3*), pLS404 (*TRP1*), pLS405 (*LEU2*), pLS406 (*URA3*), pLS408 (*natMX6*), and pLS410 (*KanMX*)^25–27^. To our knowledge, the pLS plasmids are the smallest shuttle vectors for yeast, ranging from approximately 2.6 kb to 3.5 kb in size. We have deposited all plasmids with Addgene. We anticipate that this study will streamline genetic engineering in *Saccharomyces cerevisiae*, facilitating a number of challenging applications.

## RESULTS

### Components of Minimal Shuttle Vectors

We took the DNA sequences for the prototrophic biosynthetic markers *HIS3, TRP1, LEU2*, and *URA3* from the S228C laboratory strain sequence published in the *Saccharomyces* Genome Database (SGD)^28–31^. For the drug resistance markers *natMX6* and *KanMX*, which confer resistance to nourseothricin and kanamycin, respectively, the DNA sequences were obtained from the SnapGene website^32,33^. For *natMX6* and *KanMX*, we adopted the commonly used promoter and terminator of the *TEF1* gene from *Ashbya gossypii*. These yeast selection markers, ranging from 1051 bp to 1905 bp, represent the largest elements in our plasmids.

For the bacterial selection marker *AmpR*, we used the sequence in the minimal cloning vector pUCmu, where a deletion enabled the resistance marker to utilize a shortened terminator sequence, reducing the size from 1095 bp to 947 bp^21^. Next, we incorporated the ORI, which is essential for plasmid propagation in bacterial hosts. The ORI sequence was derived from the minimal cloning vector pUCmini, where a random deletion mutation of the pUC variant of the pMB1 ORI was identified to reduce the size of the plasmid, resulting in an ORI of just 589 bp compared to 750 bp^21,34^.

The MCS spans 56 bp and includes 10 unique restriction sites for widely used 6-cutter enzymes: *BssHII, Eco53kI*/*SacI, KpnI*/*Acc65I, SmaI*/*XmaI, BamHI, XbaI, SalI, PstI* (except in pLS406, pLS408, and pLS410 due to its presence in the promoter of the yeast selection marker), *SphI, AvrII*. This MCS is similar to that of pICOz^21^. To facilitate seamless sequencing and verification of insert integration, we positioned the M13 forward and reverse sequences, each 17 bp in length, to flank the MCS.

### Introduction of Mutations to Eliminate Restriction Sites Outside MCS

We removed nearly all restriction enzyme recognition sites outside the MCS (Tables 1, 2). Because our DNA sequences were synthesized *de novo*, we could easily introduce multiple mutations in the open reading frames (ORFs) and terminators of the selection markers *AmpR, HIS3, TRP1, LEU2, URA3, natMX6*, and *KanMX* at the same time. The changes in the ORF DNA sequences exclusively switched synonymous codons. Hence, we removed all 6-bp and 8-bp restriction enzyme recognition sites (unless the restriction site was also present in a promoter) without altering the encoded amino acid sequences^35^. Furthermore, we introduced mutations in the gene terminators, substituting guanine and cytosine with adenine and thymine, to recover additional restriction sites within the selection markers. However, to prevent potential disruptions in gene expression, we did not modify promoters. Similarly, the ORI was left unchanged to ensure that plasmid replication in the bacterial host remained unaffected. On average, 68 mutations were introduced per minimal shuttle vector, effectively removing unnecessary restriction sites while preserving the original functionality. Since some of the recovered sites are still present in the regions of the plasmid backbone we did not change (promoters, ORI), the final number of recovered restriction sites is reduced. Information on the number of mutations introduced in each selection marker and recovered restriction sites per minimal shuttle vector is provided in Table 1. Detailed information on the restriction enzyme recognition sites recovered for each selectable marker is provided in Table 2. As shown in Figure 1, nearly all restriction sites were removed from the ORFs and terminators of the bacterial and yeast selection markers, leaving only a few restriction sites in the promoters and ORI regions. The final plasmid sequences have been validated through whole-plasmid sequencing, and the DNA sequences of the pLS minimal shuttle vectors are provided in Supplementary Notes 1-6 and as annotated sequence maps in gbk format in Supplementary Files.

**Table 1.** Overview of introduced mutations and recovered restriction sites. Some recoded restriction sites remain present in the unchanged regions of the plasmid backbone, that is, the promoters and ORI, reducing the number of overall recovered restriction sites.

|  | *AmpR* | *HIS3* | *TRP1* | *LEU2* | *URA3* | *natMX6* | *KanMX* |
| --- | --- | --- | --- | --- | --- | --- | --- |
| Number of introduced mutations | 36 | 28 | 20 | 43 | 34 | 37 | 31 |
| Number of recovered cut sites in gene | 33 | 24 | 15 | 27 | 26 | 29 | 17 |
| Recovered cut sites in plasmid |  | <u>pLS403</u><br>33 | <u>pLS404</u><br>24 | <u>pLS405</u><br>31 | <u>pLS406</u><br>34 | <u>pLS408</u><br>33 | <u>pLS410</u><br>27 |

**Table 2.**
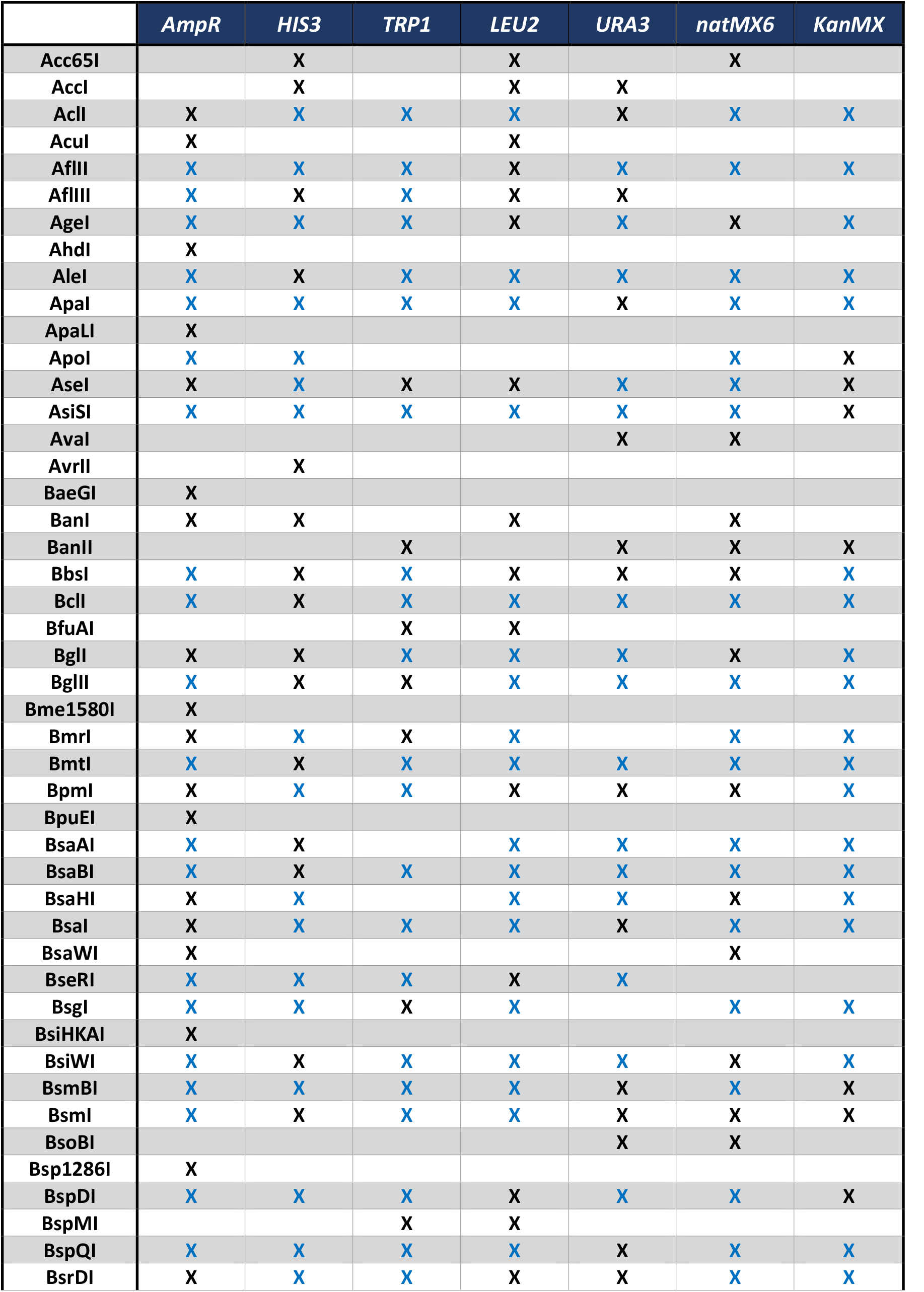
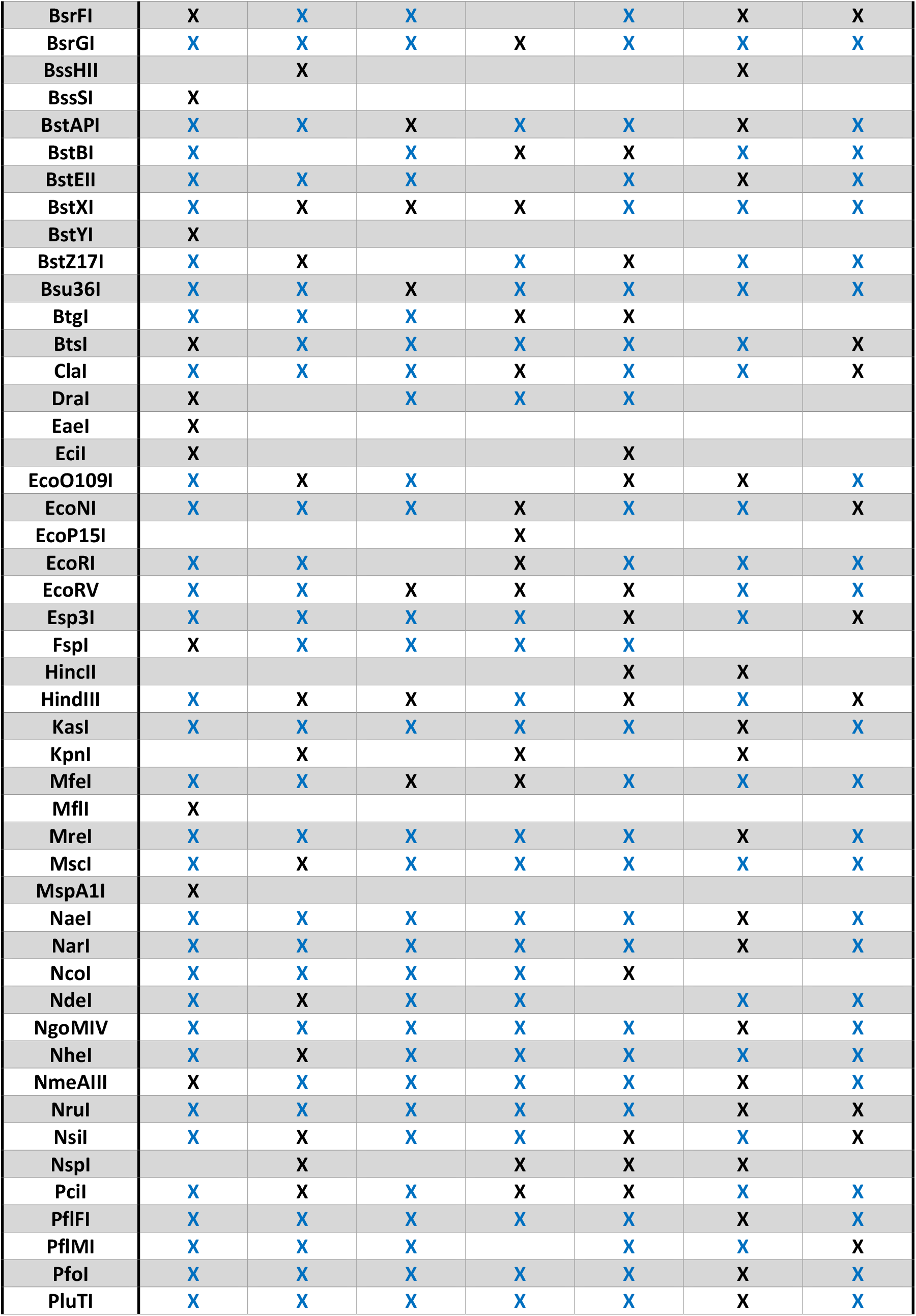
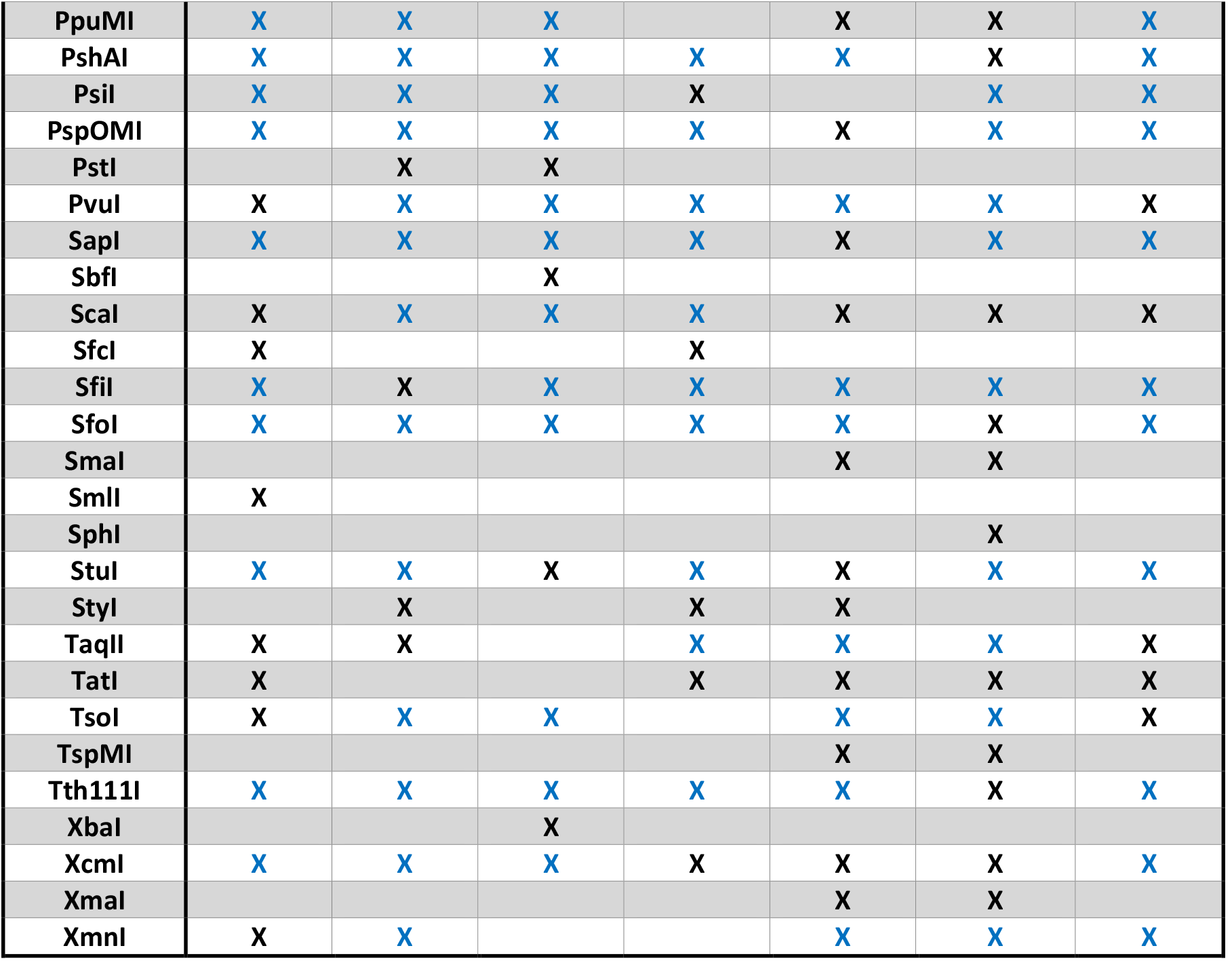
Overview of restriction sites in genes in the pLS minimal shuttle vectors. Black crosses indicate restriction sites that have been recovered; blue crosses indicate restrictions that were already absent. The table includes restriction enzymes with degenerate recognition sequences, which may result in redundancies among the recovered cut sites recognized by multiple enzymes.

### Functional Validation of the pLS Minimal Shuttle Vectors in *Saccharomyces cerevisiae*

We validated the functionality of the pLS minimal shuttle vectors for genetic manipulation in both *Escherichia coli* and *Saccharomyces cerevisiae*. The bacterial selection marker *AmpR* and the compact ORI were confirmed to be functional, as evidenced by the successful propagation of all plasmids in XL10 *Escherichia coli* cells. We observed robust bacterial growth with transformation plates exhibiting a high density of colonies and efficient DNA recovery after miniprep purification (data not shown). To validate the functionality of the recoded yeast selection markers and the utility of the minimal shuttle vectors for genetic manipulation of *Saccharomyces cerevisiae*, we introduced sequences from the non-essential gene YCR051W into the MCS of pLS403, pLS404, pLS405, pLS406, pLS408, and pLS410 for single-copy genomic integration. The ability of the pLS minimal shuttle vectors to rescue auxotrophic mutations in the haploid W303 yeast strain background confirmed their functionality (Fig. 2). Compared to their respective negative controls, the pLS vectors successfully complemented the auxotrophic deficiencies, as demonstrated by high-density colony growth on drop-out plates, validating the functionality of the recoded yeast selection markers. The DNA sequences of the single-copy integration fragment YCR051W and the primers used for its amplification are provided in Supplementary Notes 7-8. Collectively, these results demonstrate the successful design and functional validation of the pLS minimal shuttle vectors, showcasing their potential to aid genetic engineering in yeast.

**Figure 2.**
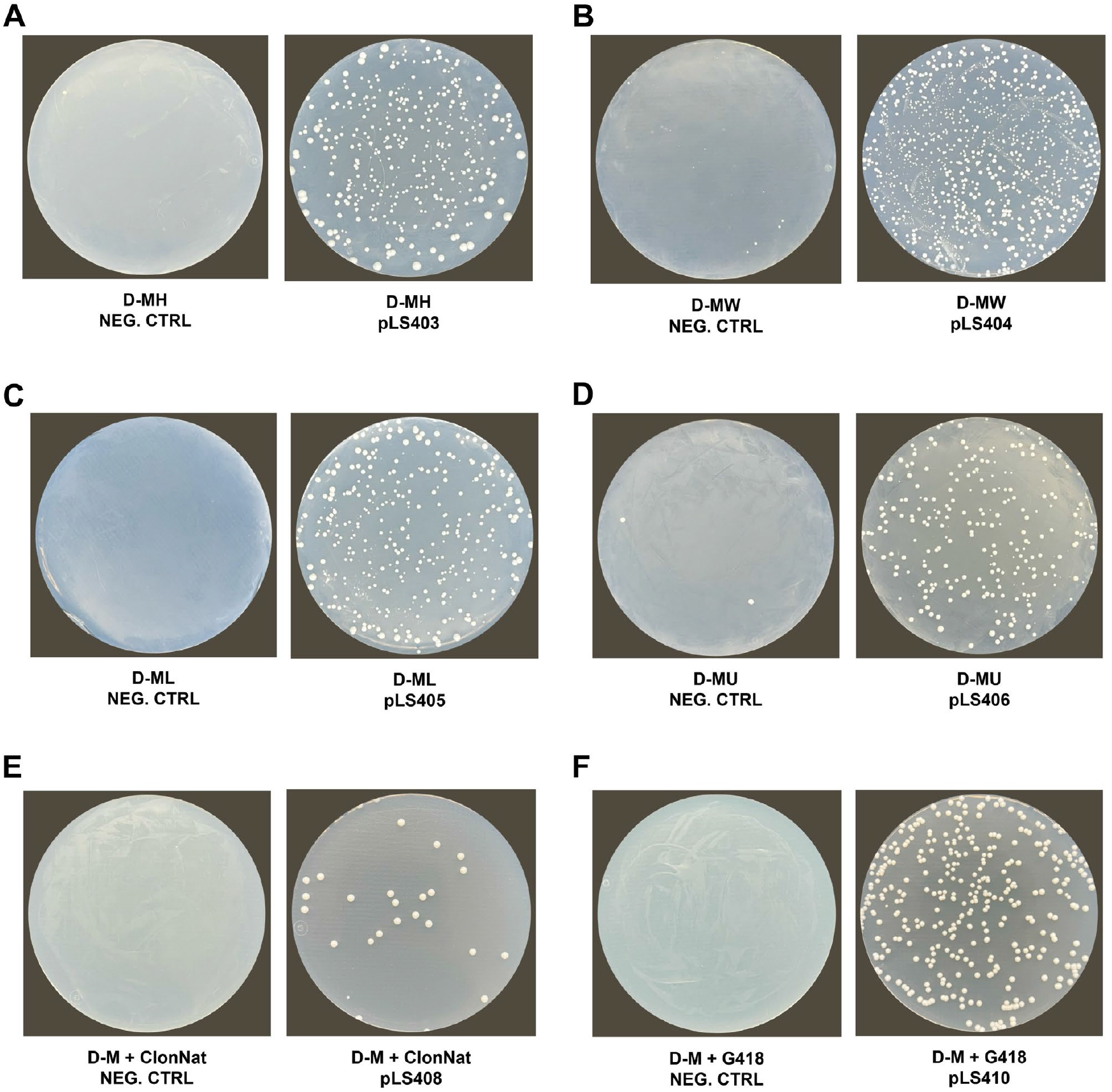
Functional validation of the pLS minimal shuttle vectors in *Saccharomyces cerevisiae*. Sequences from the non-essential gene YCR051W were inserted into the Multiple Cloning Site (MCS) of the plasmids to allow single-copy integration into the YCR051W locus. Yeast transformation of (A) pLS403, (B) pLS404, (C) pLS405, (D) pLS406, (E) pLS408, (F) pLS410. Transformed strains were selected on drop-out plates lacking histidine (D-MH), tryptophan (D-MW), leucine (D-ML), or uracil (D-MU), or on plates supplemented with the antibiotics ClonNat (D-M + ClonNat) or G418 (D-M + G418) for the selection of the *KanMX* or *natMX6* markers, respectively. All growth medium plates also lacked methionine (-M). Negative (no-DNA) controls are shown on the left.

## DISCUSSION

The design of the pLS series simplifies a range of potential applications. For example, the small size of the vectors allows the entire backbone to be amplified more easily by PCR. This simplifies mutagenesis, allowing specific mutations to be introduced into target inserts using divergent primers. The small size also facilitated the construction of the other minimal shuttle vectors starting with pLS405, as described in Materials and Methods.

Moreover, the lack of restriction sites in the pLS backbone enhances the vectors’ utility for genetic manipulation of inserts. For instance, it allows the insertion of regulatory domains at various unique restriction sites in the target. Examples of such applications are in the work by M. L. Azoitei et al. and K. A. Reynolds et al., who screened for grafting sites within the target to identify clones with optimal functional activity^36,37^.

Further optimization of the pLS vectors may be possible by replacing the selection markers with shorter alternatives, performing deletion engineering of the promoters and terminators, or by utilizing size-reduced origins of replication. For example, the minimal ORI from the low-copy plasmid pSC101 is only 220 bp in length but requires an additional initiator protein^38^. With regard to the selection markers, replacing *AmpR* with the *dfrB10* gene, which is only 237 bp long^23^, or nano-antibiotics^39,40^ may be considered.

In conclusion, our study introduces a novel set of minimal shuttle vectors tailored for yeast applications, harboring the marker genes *HIS3* (pLS403), *TRP1* (pLS404), *LEU2* (pLS405), *URA3* (pLS406), *natMX6* (pLS408), and *KanMX* (pLS410). By removing non-essential elements and strategically eliminating restriction sites, these vectors provide valuable tools for genetic manipulation in *Saccharomyces cerevisiae*.

## MATERIALS AND METHODS

### Plasmid construction

The DNA sequence of pLS405 was synthesized by GenScript. After validating the functionality of pLS405, we constructed the other minimal shuttle vectors pLS403, pLS404, pLS406, pLS408, and pLS410 through Gibson Assembly (New England Biolabs, USA). The common plasmid backbone was amplified from pLS405, while the recoded yeast selection marker sequences (*HIS3, TRP1, URA3, natMX6*, and *KanMX*) were synthesized by Twist Bioscience, and subsequently cloned into the amplified backbone. DNA digestions were performed using restriction endonucleases (New England Biolabs, USA). All PCRs were performed using high-fidelity Phusion Polymerase (New England Biolabs, USA). The DNA sequences of the constructs were verified by Sanger sequencing (Microsynth AG, Switzerland). The DNA sequences of the minimal shuttle vectors pLS403, pLS404, pLS405, pLS406, pLS408, and pLS410 are provided in Supplementary Notes 1-6.

### Bacterial strains and media

Plasmids were propagated in XL10 *Escherichia coli* cells cultured in LB medium, which was supplemented with ampicillin to select for plasmid-containing bacteria.

### Yeast strains and media

The experimental validation of the minimal shuttle vectors was conducted using the wild-type haploid W303 yeast strain background (*MATa ade2-1 leu2-3 ura3-1 trp1-1 his3-11,15 can1-100*). Yeast transformations were performed using the LiAc/DNA carrier/PEG (polyethylene glycol) protocol^41^. Transformed strains were selected on drop-out plates lacking histidine, tryptophan, leucine, or uracil, or on plates supplemented with antibiotics ClonNat or G418 for the selection of *natMX6* or *KanMX* markers, respectively. Synthetic Complete (SC) media without methionine (-Met), supplemented with 2% (w/v) glucose (D) and 0.1% (w/v) monosodium glutamate (instead of ammonium sulfate) was used for culturing^42^. For preparing solid SC plates, 2% agar and 0.17% yeast nitrogen base were added to the medium.

### Experimental validation

To validate the functionality of the recoded yeast selection markers and the usability of the minimal shuttle vectors for genetic manipulation of *Saccharomyces cerevisiae*, we inserted sequences from the non-essential gene YCR051W into the Multiple Cloning Site (MCS) of pLS403, pLS404, pLS405, pLS406, pLS408, and pLS410, which allowed single-copy integration. Cloning was performed using the restriction endonucleases BssHII and SalI (New England Biolabs, USA), while ligation was performed with T4 ligase (New England Biolabs, USA). Prior to yeast transformation, plasmid linearization was achieved using the restriction endonuclease AfeI, which is present within the YCR051W sequence. The ability of the pLS minimal shuttle vectors to rescue the auxotrophic mutations in the haploid W303 yeast strain background confirmed their functionality.

## DATA AVAILABILITY

The data generated in the paper are available from the corresponding author upon request. All plasmids and maps have been deposited with Addgene.

## CONTRIBUTIONS

L.S. constructed the plasmids, created the bacterial and yeast strains, performed the functional validation, and analyzed the data. L.S. and S.R. conceptualized the project and wrote the manuscript. S.R. and P.B. supervised the project and acquired funding.

## ACKNOWLEDGEMENTS

L.S. is supported by the EPFLglobaLeaders doctoral fellowship; an EPFL Science Seed Fund awarded to P.B. and S.J.R.; and SNSF grants CRSK-3_190526, 310030_204938, and CRSK-3_221036 awarded to S.J.R.

## Funded by

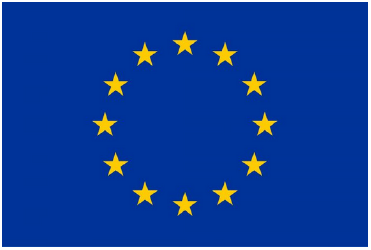

This project has received funding from the European Union’s Horizon 2020 research and innovation program under the Marie Skłodowska-Curie grant agreement No 945363.

## SUPPLEMENTARY MATERIALS

### Supplementary Note 1. DNA Sequence of pLS403

TATGTATCCGCTCATGAGACAATAACCCTGATAAATGCTTCAATAATATTGAAAAAGGAAGAGTATGAGTATTCAACATTTCCGTGTCGCCCTTATTCCCTTTTTTGCGGCATTTTGCCTTCCTGTTTTTGCTCACCCAGAAACGCTGGTGAAAGTAAAAGATGCAGAAGATCAGTTGGGTGCGCGAGTGGGTTACATCGAACTGGACCTCAACAGTGGTAAGATACTAGAGAGTTTTCGCCCCGAAGAACGATTCCCAATGATGTCCACTTTTAAGGTTCTGCTATGTGGCGCGGTATTATCCCGTATTGACGCGGGGCAAGAGCAACTAGGTCGCCGCATACACTATTCTCAGAATGACTTGGTTGAGTATTCACCAGTCACAGAAAAGCATCTTACGGATGGCATGACAGTAAGAGAATTATGTAGTGCTGCCATAACGATGAGTGATAACACGGCGGCGAACTTACTTCTGACAACGATTGGAGGACCAAAGGAGCTAACCGCTTTTTTGCACAACATGGGGGATCATGTAACTCGCCTTGATCGTTGGGAACCAGAGCTGAATGAAGCCATACCAAACGACGAGCGTGACACCACGATGCCTGTTGCTATGGCAACAACCTTGCGGAAACTATTAACTGGCGAACTACTTACTCTAGCTTCCCGGCAACAGTTAATAGACTGGATGGAGGCTGATAAAGTTGCAGGACCACTTCTGCGCTCTGCGCTTCCGGCTGGCTGGTTTATTGCTGATAAATCTGGGGCCGGAGAGCGTGGGTCACGCGGTATCATAGCAGCACTAGGGCCAGATGGTAAGCCCTCCCGTATCGTAGTTATCTACACGACGGGGAGCCAGGCAACTATGGATGAACGAAATAGACAGATCGCTGAGATAGGTGCGTCACTGATTAAGCATTGGTAGTAGAAAAGATCAAAGGATCTTCTTGAGATCCTTTTTTTCTGCGCGTAATCTGCTGCTTGCAAACAAAAAAACCACCGCTACCAGCGGTGGTTTGTTTGCCGGATCAAGAGCTACCAACTCTTTTTCCGAAGGTAACTGGCTTCAGCAGAGCGCAGATACCAAATACTGTCCTTCTAGTGTAGCCGTAGTTAGGCCACCACTTCAAGAACTCTGTAGCACCGCCTACATACCTCGCTCTGCTAATCCTGTTACCAGTGGCTGCTGCCAGTGGCGATAAGTCGTGTCTTACCGGGTTGGACTCAAGACGATAGTTACCGGATAAGGCGCAGCGGTCGGGCTGAACGGGGGGTTCGTGCACACAGCCCAGCTTGGAGCGAACGACCTACACCGAACTGAGATACCTACAGCGTGAGCTATGAGAAAGCGCCACGCTTCCCGAAGGGAGAAAGGCGGACAGGTATCCGGTAAGCGGCAGGGTCGGAACAGGAGAGCGCACGAGGGAGCTTCCAGGGGGAAACGCCTGGTATCTTTATAGTCCTGTCGGGTTTCGCCACCTCTGACTTGAGCGTCGATTTTTGTGATGCTCGTCAGGGGGGCGGAGCCTATGGAAACAGGAAACAGCTATGACGCGCGCGAGCTCGGTACCCGGGATCCTCTAGAGTCGACCTGCAGGCATGCCCTAGGACTGGCCGTCGTTTTACCTAGTACACTCTATATTTTTTTATGCCTCGGTAATGATTTTCATTTTTTTTTTTCCACCTAGCGGATGACTCTTTTTTTTTCTTAGCGATTGGCATTATCACATAATGAATTATACATTATATAAAGTAATGTGATTTCTTCGAAGAATATACTAAAAAATGAGCAGGCAAGATAAACGAAGGCAAAGATGACAGAGCAGAAAGCCCTAGTAAAGCGTATTACAAATGAAACCAAGATTCAGATTGCGATTTCTTTAAAGGGTGGTCCCCTAGCGATAGAACACTCGATCTTCCCAGAAAAAGAGGCAGAAGCAGTAGCAGAACAGGCCACACAATCGCAAGTGATTAACGTCCACACAGGTATAGGGTTTCTGGACCACATGATTCATGCTCTCGCCAAGCACTCCGGCTGGTCGCTAATCGTTGAGTGCATTGGTGACTTACACATAGACGACCATCACACCACTGAGGACTGCGGGATTGCTCTGGGTCAGGCTTTTAAAGAGGCTCTAGGGGCCGTGCGTGGAGTAAAAAGGTTTGGATCAGGATTTGCGCCTTTGGATGAGGCACTTTCCAGGGCGGTTGTTGATCTTTCGAACAGGCCATACGCAGTTGTCGAACTTGGTTTGCAAAGGGAGAAAGTAGGTGATCTCTCTTGCGAGATGATCCCGCATTTTCTTGAAAGTTTTGCAGAGGCAAGCAGAATTACCCTCCACGTTGATTGTCTGCGAGGCAAGAATGACCATCACCGTAGCGAGAGTGCGTTCAAGGCTCTTGCGGTTGCCATAAGAGAAGCCACCTCGCCCAATGGAACCAACGATGTTCCCTCCACCAAAGGTGTTCTTATGTAGTGACACCGATTATTTAAAGCTGCATCATACGATATATATATATGTGTATATATGTATATCTATGAATGTCAGTAAGTATGTATATGAACAGTATGATACTGAAAATGACAAGGTAATACATCATTCTATATGTGTCATTCTGAACGAGGCGCGTTTTCCTTTTTTCTTTTTGCTTTTTCTTTTTTTTTCTCTTGAACTCG

### Supplementary Note 2. DNA Sequence of pLS404

TATGTATCCGCTCATGAGACAATAACCCTGATAAATGCTTCAATAATATTGAAAAAGGAAGAGTATGAGTATTCAACATTTCCGTGTCGCCCTTATTCCCTTTTTTGCGGCATTTTGCCTTCCTGTTTTTGCTCACCCAGAAACGCTGGTGAAAGTAAAAGATGCAGAAGATCAGTTGGGTGCGCGAGTGGGTTACATCGAACTGGACCTCAACAGTGGTAAGATACTAGAGAGTTTTCGCCCCGAAGAACGATTCCCAATGATGTCCACTTTTAAGGTTCTGCTATGTGGCGCGGTATTATCCCGTATTGACGCGGGGCAAGAGCAACTAGGTCGCCGCATACACTATTCTCAGAATGACTTGGTTGAGTATTCACCAGTCACAGAAAAGCATCTTACGGATGGCATGACAGTAAGAGAATTATGTAGTGCTGCCATAACGATGAGTGATAACACGGCGGCGAACTTACTTCTGACAACGATTGGAGGACCAAAGGAGCTAACCGCTTTTTTGCACAACATGGGGGATCATGTAACTCGCCTTGATCGTTGGGAACCAGAGCTGAATGAAGCCATACCAAACGACGAGCGTGACACCACGATGCCTGTTGCTATGGCAACAACCTTGCGGAAACTATTAACTGGCGAACTACTTACTCTAGCTTCCCGGCAACAGTTAATAGACTGGATGGAGGCTGATAAAGTTGCAGGACCACTTCTGCGCTCTGCGCTTCCGGCTGGCTGGTTTATTGCTGATAAATCTGGGGCCGGAGAGCGTGGGTCACGCGGTATCATAGCAGCACTAGGGCCAGATGGTAAGCCCTCCCGTATCGTAGTTATCTACACGACGGGGAGCCAGGCAACTATGGATGAACGAAATAGACAGATCGCTGAGATAGGTGCGTCACTGATTAAGCATTGGTAGTAGAAAAGATCAAAGGATCTTCTTGAGATCCTTTTTTTCTGCGCGTAATCTGCTGCTTGCAAACAAAAAAACCACCGCTACCAGCGGTGGTTTGTTTGCCGGATCAAGAGCTACCAACTCTTTTTCCGAAGGTAACTGGCTTCAGCAGAGCGCAGATACCAAATACTGTCCTTCTAGTGTAGCCGTAGTTAGGCCACCACTTCAAGAACTCTGTAGCACCGCCTACATACCTCGCTCTGCTAATCCTGTTACCAGTGGCTGCTGCCAGTGGCGATAAGTCGTGTCTTACCGGGTTGGACTCAAGACGATAGTTACCGGATAAGGCGCAGCGGTCGGGCTGAACGGGGGGTTCGTGCACACAGCCCAGCTTGGAGCGAACGACCTACACCGAACTGAGATACCTACAGCGTGAGCTATGAGAAAGCGCCACGCTTCCCGAAGGGAGAAAGGCGGACAGGTATCCGGTAAGCGGCAGGGTCGGAACAGGAGAGCGCACGAGGGAGCTTCCAGGGGGAAACGCCTGGTATCTTTATAGTCCTGTCGGGTTTCGCCACCTCTGACTTGAGCGTCGATTTTTGTGATGCTCGTCAGGGGGGCGGAGCCTATGGAAACAGGAAACAGCTATGACGCGCGCGAGCTCGGTACCCGGGATCCTCTAGAGTCGACCTGCAGGCATGCCCTAGGACTGGCCGTCGTTTTACAACGACATTACTATATATATAATATAGGAAGCATTTAATAGAACAGCATCGTAATATATGTGTACTTTGCAGTTATGACGCCAGATGGCAGTAGTGGAAGATATTCTTTATTGAAAAATAGCTTGTCACCTTACGTACAATCTTGATCCGGAGCTTTTCTTTTTTTGCCGATTAAGAATTCGGTCGAAAAAAGAAAAGGAGAGGGCCAAGAGGGAGGGCATTGGTGACTATTGAGCACGTGAGTATACGTGATTAAGCACACAAAGGCAGCTTGGAGTATGTCTGTTATCAATTTCACAGGTAGTTCTGGTCCATTGGTGAAAGTTTGCGGCTTGCAGAGCACAGAGGCCGCCGAATGTGCTCTTGATTCCGATGCTGACTTGCTCGGTATTATATGTGTCCCCAATAGAAAGAGAACTATTGACCCGGTTATTGCAAGGAAAATTTCAAGTCTTGTAAAAGCATATAAAAATAGTTCAGGCACTCCGAAATACTTGGTTGGCGTGTTTCGTAATCAACCAAAGGAGGATGTTTTGGCTCTGGTCAATGATTACGGCATTGATATAGTGCAACTGCATGGTGACGAGTCGTGGCAAGAATACCAAGAGTTCCTCGGTTTGCCAGTTATTAAAAGACTCGTATTTCCAAAAGACTGCAACATACTACTCAGCGCAGCTTCACAGAAACCTCATTCGTTTATTCCCTTGTTTGATTCAGAAGCGGGTGGGACAGGTGAACTTTTGGATTGGAACTCGATTTCTGATTGGGTAGGAAGGCAAGAGAGTCCCGAAAGTTTACATTTTATGTTAGCTGGTGGACTGACGCCAGAAAATGTTGGTGATGCGCTTAGATTAAATGGCGTTATTGGTGTTGATGTAAGCGGAGGTGTGGAGACAAATGGTGTAAAAGACTCTAACAAAATAGCAAATTTCGTCAAAAATGCTAAGAAATAGGTTATTACTGAGTAGTATTTATTTAAGTATTGTTTGTGCATTTGCCTGCAAGCCTTTTGAAAAGCAAGCATAAAATATCTAAACATAAAATCTGTAAAAT

### Supplementary Note 3. DNA Sequence of pLS405

TATGTATCCGCTCATGAGACAATAACCCTGATAAATGCTTCAATAATATTGAAAAAGGAAGAGTATGAGTATTCAACATTTCCGTGTCGCCCTTATTCCCTTTTTTGCGGCATTTTGCCTTCCTGTTTTTGCTCACCCAGAAACGCTGGTGAAAGTAAAAGATGCAGAAGATCAGTTGGGTGCGCGAGTGGGTTACATCGAACTGGACCTCAACAGTGGTAAGATACTAGAGAGTTTTCGCCCCGAAGAACGATTCCCAATGATGTCCACTTTTAAGGTTCTGCTATGTGGCGCGGTATTATCCCGTATTGACGCGGGGCAAGAGCAACTAGGTCGCCGCATACACTATTCTCAGAATGACTTGGTTGAGTATTCACCAGTCACAGAAAAGCATCTTACGGATGGCATGACAGTAAGAGAATTATGTAGTGCTGCCATAACGATGAGTGATAACACGGCGGCGAACTTACTTCTGACAACGATTGGAGGACCAAAGGAGCTAACCGCTTTTTTGCACAACATGGGGGATCATGTAACTCGCCTTGATCGTTGGGAACCAGAGCTGAATGAAGCCATACCAAACGACGAGCGTGACACCACGATGCCTGTTGCTATGGCAACAACCTTGCGGAAACTATTAACTGGCGAACTACTTACTCTAGCTTCCCGGCAACAGTTAATAGACTGGATGGAGGCTGATAAAGTTGCAGGACCACTTCTGCGCTCTGCGCTTCCGGCTGGCTGGTTTATTGCTGATAAATCTGGGGCCGGAGAGCGTGGGTCACGCGGTATCATAGCAGCACTAGGGCCAGATGGTAAGCCCTCCCGTATCGTAGTTATCTACACGACGGGGAGCCAGGCAACTATGGATGAACGAAATAGACAGATCGCTGAGATAGGTGCGTCACTGATTAAGCATTGGTAGTAGAAAAGATCAAAGGATCTTCTTGAGATCCTTTTTTTCTGCGCGTAATCTGCTGCTTGCAAACAAAAAAACCACCGCTACCAGCGGTGGTTTGTTTGCCGGATCAAGAGCTACCAACTCTTTTTCCGAAGGTAACTGGCTTCAGCAGAGCGCAGATACCAAATACTGTCCTTCTAGTGTAGCCGTAGTTAGGCCACCACTTCAAGAACTCTGTAGCACCGCCTACATACCTCGCTCTGCTAATCCTGTTACCAGTGGCTGCTGCCAGTGGCGATAAGTCGTGTCTTACCGGGTTGGACTCAAGACGATAGTTACCGGATAAGGCGCAGCGGTCGGGCTGAACGGGGGGTTCGTGCACACAGCCCAGCTTGGAGCGAACGACCTACACCGAACTGAGATACCTACAGCGTGAGCTATGAGAAAGCGCCACGCTTCCCGAAGGGAGAAAGGCGGACAGGTATCCGGTAAGCGGCAGGGTCGGAACAGGAGAGCGCACGAGGGAGCTTCCAGGGGGAAACGCCTGGTATCTTTATAGTCCTGTCGGGTTTCGCCACCTCTGACTTGAGCGTCGATTTTTGTGATGCTCGTCAGGGGGGCGGAGCCTATGGAAACAGGAAACAGCTATGACGCGCGCGAGCTCGGTACCCGGGATCCTCTAGAGTCGACCTGCAGGCATGCCCTAGGACTGGCCGTCGTTTTACAACTGTGGGAATACTCAGGTATCGTAAGATGCAAGAGTTCGAGTCTCTTAGCAACCATTATTTTTTTCCTCAACATAACGAGAACACACAGGGGCGCTATCGCACAGAATCAAATTCGATGACTGGAAATTTTTTGTTAATTTCAGAGGTCGCCTGACGCATATACCTTTTTCAACTGAAAAATTGGGAGAAAAAGGAAAGGTGAGAGGCCGGAACCGGCTTTTCATATAGAATAGAGAAGCGTTCATGACTAAATGCTTGCATCACAATACTTGAAGTTGACAATATTATTTAAGGACCTATTGTTTTTTCCAATAGGTGGTTAGCAATCGTCTTACTTTCTAACTTTTCTTACCTTTTACATTTCAGCAATATATATATATATTTCAAGGATATACCATTCTAATGTCTGCCCCTAAGAAGATCGTCGTTTTGCCAGGGGACCACGTTGGTCAAGAAATCACAGCCGAAGCCATTAAGGTTCTTAAAGCTATTTCTGATGTTCGTTCCAATGTCAAGTTCGATTTCGAGAATCATTTAATTGGTGGTGCTGCTATCGACGCTACTGGTGTCCCACTTCCAGATGAGGCTCTGGAAGCCTCCAAGAAGGCTGATGCCGTTTTGTTAGGTGCTGTGGGTGGTCCTAAATGGGGTACGGGTAGTGTTAGACCAGAACAAGGTTTACTAAAAATCCGTAAAGAACTTCAATTATACGCCAATTTAAGACCATGTAACTTTGCATCCGATTCTCTTTTAGACTTATCTCCAATCAAGCCACAATTTGCTAAAGGTACTGACTTCGTTGTTGTCAGAGAATTAGTGGGAGGTATTTACTTTGGTAAGAGAAAGGAGGACGATGGTGATGGTGTCGCTTGGGATAGTGAACAATACACCGTTCCAGAAGTGCAAAGAATCACAAGAATGGCGGCTTTCATGGCCCTACAACATGAGCCACCTTTACCTATTTGGTCCTTAGATAAAGCTAATCTTTTGGCCTCATCAAGATTATGGAGAAAAACTGTGGAGGAAACCATCAAGAACGAGTTTCCTACATTGAAGGTTCAACATCAGTTGATTGATTCTGCCGCCATGATCCTAGTTAAGAACCCAACCCACCTAAACGGTATTATTATCACCAGCAATATGTTTGGTGATATTATCTCCGATGAAGCCTCCGTTATCCCAGGTTCGTTGGGTTTGTTGCCATCTGCGTCGTTGGCCTCTTTGCCAGACAAGAACACCGCATTTGGTTTGTACGAACCATGTCACGGTTCTGCGCCAGATTTGCCAAAGAATAAGGTTGATCCTATCGCCACTATCTTGTCTGCGGCTATGATGTTGAAATTGTCATTGAACTTGCCAGAAGAAGGTAAGGCCATTGAAGATGCAGTTAAAAAGGTTTTGGATGCAGGAATCAGAACTGGTGATTTAGGTGGTTCCAATAGTACCACCGAAGTCGGTGATGCTGTCGCCGAAGAAGTTAAGAAAATCCTTGCTTAAAAAGATTCTCTTTTTTTATGATATTTATACATAAACTATATAAATGAAATTCATAATAGAAACGACACGAAATTACAAAATGGAATATGTTCATAGGATATACGAAACTATATACGCAATCTACATACATTTATCAAGAAGGAGAAAAAGGAGGATAGTAAAGGAATACAGGTAAGCAAATTGATACTAATGGCTCAACGTGATAAGGAAAAAGAATTGCACTTTAACATTAAAATTGACAAGGAATAGGTCACCACACAAAAAGTTAGGTGTAACAGAAAATCATAAAACTACGATTCCTAATTTGATATTGGAGGATTTTCTCTAAAAAAAAAAAAATACAACAAATAAAAAACACTCAATGACCTGACCATTTGATGGAGTTTAAGTCAATACCTTCTTGAA

### Supplementary Note 4. DNA Sequence of pLS406

TATGTATCCGCTCATGAGACAATAACCCTGATAAATGCTTCAATAATATTGAAAAAGGAAGAGTATGAGTATTCAACATTTCCGTGTCGCCCTTATTCCCTTTTTTGCGGCATTTTGCCTTCCTGTTTTTGCTCACCCAGAAACGCTGGTGAAAGTAAAAGATGCAGAAGATCAGTTGGGTGCGCGAGTGGGTTACATCGAACTGGACCTCAACAGTGGTAAGATACTAGAGAGTTTTCGCCCCGAAGAACGATTCCCAATGATGTCCACTTTTAAGGTTCTGCTATGTGGCGCGGTATTATCCCGTATTGACGCGGGGCAAGAGCAACTAGGTCGCCGCATACACTATTCTCAGAATGACTTGGTTGAGTATTCACCAGTCACAGAAAAGCATCTTACGGATGGCATGACAGTAAGAGAATTATGTAGTGCTGCCATAACGATGAGTGATAACACGGCGGCGAACTTACTTCTGACAACGATTGGAGGACCAAAGGAGCTAACCGCTTTTTTGCACAACATGGGGGATCATGTAACTCGCCTTGATCGTTGGGAACCAGAGCTGAATGAAGCCATACCAAACGACGAGCGTGACACCACGATGCCTGTTGCTATGGCAACAACCTTGCGGAAACTATTAACTGGCGAACTACTTACTCTAGCTTCCCGGCAACAGTTAATAGACTGGATGGAGGCTGATAAAGTTGCAGGACCACTTCTGCGCTCTGCGCTTCCGGCTGGCTGGTTTATTGCTGATAAATCTGGGGCCGGAGAGCGTGGGTCACGCGGTATCATAGCAGCACTAGGGCCAGATGGTAAGCCCTCCCGTATCGTAGTTATCTACACGACGGGGAGCCAGGCAACTATGGATGAACGAAATAGACAGATCGCTGAGATAGGTGCGTCACTGATTAAGCATTGGTAGTAGAAAAGATCAAAGGATCTTCTTGAGATCCTTTTTTTCTGCGCGTAATCTGCTGCTTGCAAACAAAAAAACCACCGCTACCAGCGGTGGTTTGTTTGCCGGATCAAGAGCTACCAACTCTTTTTCCGAAGGTAACTGGCTTCAGCAGAGCGCAGATACCAAATACTGTCCTTCTAGTGTAGCCGTAGTTAGGCCACCACTTCAAGAACTCTGTAGCACCGCCTACATACCTCGCTCTGCTAATCCTGTTACCAGTGGCTGCTGCCAGTGGCGATAAGTCGTGTCTTACCGGGTTGGACTCAAGACGATAGTTACCGGATAAGGCGCAGCGGTCGGGCTGAACGGGGGGTTCGTGCACACAGCCCAGCTTGGAGCGAACGACCTACACCGAACTGAGATACCTACAGCGTGAGCTATGAGAAAGCGCCACGCTTCCCGAAGGGAGAAAGGCGGACAGGTATCCGGTAAGCGGCAGGGTCGGAACAGGAGAGCGCACGAGGGAGCTTCCAGGGGGAAACGCCTGGTATCTTTATAGTCCTGTCGGGTTTCGCCACCTCTGACTTGAGCGTCGATTTTTGTGATGCTCGTCAGGGGGGCGGAGCCTATGGAAACAGGAAACAGCTATGACGCGCGCGAGCTCGGTACCCGGGATCCTCTAGAGTCGACCTGCAGGCATGCCCTAGGACTGGCCGTCGTTTTACTTCAATTCATCATTTTTTTTTTATTCTTTTTTTTGATTTCGGTTTCCTTGAAATTTTTTTGATTCGGTAATCTCCGAACAGAAGGAAGAACGAAGGAAGGAGCACAGACTTAGATTGGTATATATACGCATATGTAGTGTTGAAGAAACATGAAATTGCCCAGTATTCTTAACCCAACTGCACAGAACAAAAACCTGCAGGAAACGAAGATAAATCATGTCGAAAGCTACATATAAGGAACGTGCTGCTACTCATCCTAGTCCTGTTGCTGCCAAGCTATTTAATATCATGCACGAAAAGCAAACAAACTTGTGTGCTTCATTGGATGTTCGTACCACGAAGGAATTACTAGAGTTAGTTGAAGCATTAGGTCCGAAAATTTGTTTACTAAAAACTCATGTGGATATTTTGACTGATTTTTCGATGGAGGGGACAGTTAAGCCGCTAAAGGCATTATCAGCCAAGTATAATTTTTTACTCTTCGAGGACAGAAAATTTGCTGACATTGGTAATACAGTCAAATTGCAGTATTCTGCGGGTGTATATAGAATAGCAGAATGGGCAGACATTACGAACGCACACGGTGTGGTGGGGCCAGGTATTGTTAGCGGTTTGAAGCAGGCGGCAGAAGAAGTAACAAAGGAACCTAGAGGGCTTTTGATGTTAGCAGAATTGTCATGCAAGGGTTCCCTAAGCACTGGTGAATATACTAAGGGTACTGTTGATATTGCCAAGAGCGACAAAGATTTTGTTATCGGCTTTATTGCTCAAAGAGACATGGGTGGAAGAGATGAAGGTTACGATTGGTTGATTATGACACCCGGTGTGGGTTTAGATGACAAGGGTGACGCATTGGGCCAACAGTATAGAACCGTAGATGATGTAGTCTCTACAGGATCAGACATTATTATTGTAGGAAGAGGACTATTTGCAAAGGGAAGGGATGCAAAGGTAGAGGGTGAGCGTTACAGAAAAGCAGGGTGGGAAGCATATTTGAGAAGATGCGGGCAGCAAAACTAAAAAACTGTATTATAAGTAAATGAATGTATAATAAACTCACAAATTAGAGCTTCAATTTAATTATATCAGTTATTACCCGGAAATCTTGGTCGTAATGATTTCTATAATGACGAAAAAAAAAAAATTGGAAAGAAAAAACTTCATGGCCTTTATAAAAAGGAACTATCCAATACCTCGCCAGAACCAAGTAACAGTATT

### Supplementary Note 5. DNA Sequence of pLS408

TATGTATCCGCTCATGAGACAATAACCCTGATAAATGCTTCAATAATATTGAAAAAGGAAGAGTATGAGTATTCAACATTTCCGTGTCGCCCTTATTCCCTTTTTTGCGGCATTTTGCCTTCCTGTTTTTGCTCACCCAGAAACGCTGGTGAAAGTAAAAGATGCAGAAGATCAGTTGGGTGCGCGAGTGGGTTACATCGAACTGGACCTCAACAGTGGTAAGATACTAGAGAGTTTTCGCCCCGAAGAACGATTCCCAATGATGTCCACTTTTAAGGTTCTGCTATGTGGCGCGGTATTATCCCGTATTGACGCGGGGCAAGAGCAACTAGGTCGCCGCATACACTATTCTCAGAATGACTTGGTTGAGTATTCACCAGTCACAGAAAAGCATCTTACGGATGGCATGACAGTAAGAGAATTATGTAGTGCTGCCATAACGATGAGTGATAACACGGCGGCGAACTTACTTCTGACAACGATTGGAGGACCAAAGGAGCTAACCGCTTTTTTGCACAACATGGGGGATCATGTAACTCGCCTTGATCGTTGGGAACCAGAGCTGAATGAAGCCATACCAAACGACGAGCGTGACACCACGATGCCTGTTGCTATGGCAACAACCTTGCGGAAACTATTAACTGGCGAACTACTTACTCTAGCTTCCCGGCAACAGTTAATAGACTGGATGGAGGCTGATAAAGTTGCAGGACCACTTCTGCGCTCTGCGCTTCCGGCTGGCTGGTTTATTGCTGATAAATCTGGGGCCGGAGAGCGTGGGTCACGCGGTATCATAGCAGCACTAGGGCCAGATGGTAAGCCCTCCCGTATCGTAGTTATCTACACGACGGGGAGCCAGGCAACTATGGATGAACGAAATAGACAGATCGCTGAGATAGGTGCGTCACTGATTAAGCATTGGTAGTAGAAAAGATCAAAGGATCTTCTTGAGATCCTTTTTTTCTGCGCGTAATCTGCTGCTTGCAAACAAAAAAACCACCGCTACCAGCGGTGGTTTGTTTGCCGGATCAAGAGCTACCAACTCTTTTTCCGAAGGTAACTGGCTTCAGCAGAGCGCAGATACCAAATACTGTCCTTCTAGTGTAGCCGTAGTTAGGCCACCACTTCAAGAACTCTGTAGCACCGCCTACATACCTCGCTCTGCTAATCCTGTTACCAGTGGCTGCTGCCAGTGGCGATAAGTCGTGTCTTACCGGGTTGGACTCAAGACGATAGTTACCGGATAAGGCGCAGCGGTCGGGCTGAACGGGGGGTTCGTGCACACAGCCCAGCTTGGAGCGAACGACCTACACCGAACTGAGATACCTACAGCGTGAGCTATGAGAAAGCGCCACGCTTCCCGAAGGGAGAAAGGCGGACAGGTATCCGGTAAGCGGCAGGGTCGGAACAGGAGAGCGCACGAGGGAGCTTCCAGGGGGAAACGCCTGGTATCTTTATAGTCCTGTCGGGTTTCGCCACCTCTGACTTGAGCGTCGATTTTTGTGATGCTCGTCAGGGGGGCGGAGCCTATGGAAACAGGAAACAGCTATGACGCGCGCGAGCTCGGTACCCGGGATCCTCTAGAGTCGACCTGCAGGCATGCCCTAGGACTGGCCGTCGTTTTACGACATGGAGGCCCAGAATACCCTCCTTGACAGTCTTGACGTGCGCAGCTCAGGGGCATGATGTGACTGTCGCCCGTACATTTAGCCCATACATCCCCATGTATAATCATTTGCATCCATACATTTTGATGGCCGCACGGCGCGAAGCAAAAATTACGGCTCCTCGCTGCAGACCTGCGAGCAGGGAAACGCTCCCCTCACAGACGCGTTGAATTGTCCCCACGCCGCGCCCCTGTAGAGAAATATAAAAGGTTAGGATTTGCCACTGAGGTTCTTCTTTCATATACTTCCTTTTAAAATCTTGCTAGGATACAGTTCTCACATCACATCCGAACATAAACAACCATGGGAACCACTCTTGACGACACGGCTTACCGCTACCGCACCAGTGTTCCGGGGGACGCAGAGGCAATCGAGGCACTGGATGGTTCCTTCACCACCGACACCGTATTCCGCGTCACCGCCACCGGGGACGGCTTCACCCTGCGGGAAGTGCCAGTGGACCCGCCCCTGACCAAAGTGTTCCCCGACGACGAATCGGACGACGAATCGGACGACGGGGAGGACGGCGACCCGGACTCACGGACGTTCGTAGCGTATGGGGACGACGGCGACCTGGCGGGCTTCGTGGTCATCTCGTACTCAGCGTGGAACCGACGGCTGACCGTCGAGGACATCGAGGTAGCCCCGGAACACCGGGGTCACGGGGTCGGGCGAGCGTTGATGGGGCTGGCGACGGAGTTCGCAGGCGAACGGGGGGCTGGTCATCTCTGGCTGGAAGTCACCAACGTAAACGCACCCGCGATCCACGCGTACCGACGAATGGGGTTCACCCTCTGCGGCCTGGACACCGCCCTGTACGACGGGACCGCCTCGGACGGCGAACGGCAGGCACTCTACATGAGTATGCCCTGCCCCTAATCAATACTGACAATAAAAAGATTCTTGTTTTCAAGAACTTGTCATTTGTATAGTTTTTTTATATTGTAGTTGTTCTATTTTAATCAAATGTTAGCGTGATTTATATTTTTTTTCGCCTCGACATCATCTGCCCAGATGCGAAGTTAAGTGCGCAGAAAGTAATATCATGCGTCAATCGTATGTGAATGATGGTCGCTATACTG

### Supplementary Note 6. DNA Sequence of pLS410

TATGTATCCGCTCATGAGACAATAACCCTGATAAATGCTTCAATAATATTGAAAAAGGAAGAGTATGAGTATTCAACATTTCCGTGTCGCCCTTATTCCCTTTTTTGCGGCATTTTGCCTTCCTGTTTTTGCTCACCCAGAAACGCTGGTGAAAGTAAAAGATGCAGAAGATCAGTTGGGTGCGCGAGTGGGTTACATCGAACTGGACCTCAACAGTGGTAAGATACTAGAGAGTTTTCGCCCCGAAGAACGATTCCCAATGATGTCCACTTTTAAGGTTCTGCTATGTGGCGCGGTATTATCCCGTATTGACGCGGGGCAAGAGCAACTAGGTCGCCGCATACACTATTCTCAGAATGACTTGGTTGAGTATTCACCAGTCACAGAAAAGCATCTTACGGATGGCATGACAGTAAGAGAATTATGTAGTGCTGCCATAACGATGAGTGATAACACGGCGGCGAACTTACTTCTGACAACGATTGGAGGACCAAAGGAGCTAACCGCTTTTTTGCACAACATGGGGGATCATGTAACTCGCCTTGATCGTTGGGAACCAGAGCTGAATGAAGCCATACCAAACGACGAGCGTGACACCACGATGCCTGTTGCTATGGCAACAACCTTGCGGAAACTATTAACTGGCGAACTACTTACTCTAGCTTCCCGGCAACAGTTAATAGACTGGATGGAGGCTGATAAAGTTGCAGGACCACTTCTGCGCTCTGCGCTTCCGGCTGGCTGGTTTATTGCTGATAAATCTGGGGCCGGAGAGCGTGGGTCACGCGGTATCATAGCAGCACTAGGGCCAGATGGTAAGCCCTCCCGTATCGTAGTTATCTACACGACGGGGAGCCAGGCAACTATGGATGAACGAAATAGACAGATCGCTGAGATAGGTGCGTCACTGATTAAGCATTGGTAGTAGAAAAGATCAAAGGATCTTCTTGAGATCCTTTTTTTCTGCGCGTAATCTGCTGCTTGCAAACAAAAAAACCACCGCTACCAGCGGTGGTTTGTTTGCCGGATCAAGAGCTACCAACTCTTTTTCCGAAGGTAACTGGCTTCAGCAGAGCGCAGATACCAAATACTGTCCTTCTAGTGTAGCCGTAGTTAGGCCACCACTTCAAGAACTCTGTAGCACCGCCTACATACCTCGCTCTGCTAATCCTGTTACCAGTGGCTGCTGCCAGTGGCGATAAGTCGTGTCTTACCGGGTTGGACTCAAGACGATAGTTACCGGATAAGGCGCAGCGGTCGGGCTGAACGGGGGGTTCGTGCACACAGCCCAGCTTGGAGCGAACGACCTACACCGAACTGAGATACCTACAGCGTGAGCTATGAGAAAGCGCCACGCTTCCCGAAGGGAGAAAGGCGGACAGGTATCCGGTAAGCGGCAGGGTCGGAACAGGAGAGCGCACGAGGGAGCTTCCAGGGGGAAACGCCTGGTATCTTTATAGTCCTGTCGGGTTTCGCCACCTCTGACTTGAGCGTCGATTTTTGTGATGCTCGTCAGGGGGGCGGAGCCTATGGAAACAGGAAACAGCTATGACGCGCGCGAGCTCGGTACCCGGGATCCTCTAGAGTCGACCTGCAGGCATGCCCTAGGACTGGCCGTCGTTTTACGACATGGAGGCCCAGAATACCCTCCTTGACAGTCTTGACGTGCGCAGCTCAGGGGCATGATGTGACTGTCGCCCGTACATTTAGCCCATACATCCCCATGTATAATCATTTGCATCCATACATTTTGATGGCCGCACGGCGCGAAGCAAAAATTACGGCTCCTCGCTGCAGACCTGCGAGCAGGGAAACGCTCCCCTCACAGACGCGTTGAATTGTCCCCACGCCGCGCCCCTGTAGAGAAATATAAAAGGTTAGGATTTGCCACTGAGGTTCTTCTTTCATATACTTCCTTTTAAAATCTTGCTAGGATACAGTTCTCACATCACATCCGAACATAAACAACCATGGGTAAGGAAAAGACTCACGTTTCGAGGCCGCGATTAAACTCCAATATGGATGCTGATTTATATGGGTATAAATGGGCTAGGGATAATGTCGGGCAATCAGGTGCGACAATCTACCGATTGTATGGGAAGCCCGATGCGCCAGAGTTGTTTCTGAAACATGGCAAAGGTAGCGTTGCCAATGATGTTACAGATGAGATGGTCAGACTAAACTGGCTGACGGAGTTTATGCCTCTACCGACCATCAAGCATTTTATCCGTACTCCTGATGACGCATGGTTGCTCACCACAGCGATCCCCGGCAAAACAGCTTTCCAGGTATTAGAAGAATATCCAGATTCAGGTGAAAACATTGTTGATGCGCTGGCTGTGTTCCTGCGTCGGTTGCACTCGATTCCTGTTTGTAATTGTCCTTTTAACAGTGATCGCGTATTTCGACTCGCACAGGCGCAATCACGAATGAATAACGGTTTGGTTGATGCGAGTGATTTTGATGACGAGCGTAATGGCTGGCCTGTTGAACAAGTCTGGAAAGAAATGCACAAACTTTTGCCATTCTCACCAGATTCAGTCGTCACTCATGGTGATTTCTCACTTGATAACCTTATTTTTGACGAGGGGAAATTGATAGGTTGTATTGATGTAGGACGAGTAGGAATCGCAGACCGCTACCAAGACCTTGCAATCCTATGGAACTGCCTCGGTGAGTTTTCTCCTTCATTACAGAAACGGCTTTTTCAAAAATATGGTATTGATAATCCTGATATGAATAAATTGCAGTTTCATTTGATGCTCGATGAGTTTTTCTAATCAATACTGACAATAAAAAGATTCTTGTTTTCAAGAACTTGTCATTTGTATAGTTTTTTTATATTGTAGTTGTTCTATTTTAATCAAATGTTAGCGTGATTTATATTTTTTTTCGCCTCGACATCATCTGCCCAGATGCGAAGTTAAGTGCGCAGAAAGTAATATCATGCGTCAATCGTATGTGAATGATGGTCGCTATACTG

### Supplementary Note 7. DNA Sequence of YCR051W cloned into the MCS for single-copy integration

TAATGTGTTGGACAACGACGGCGATACCCCGTTGCACCATGTGGAGGATGTGGCCACTGCCAGGTTGATCGTGGAAGAGCTGGGTGGAGACTTCACTATCAGGAATGTGGAGGGCCAAACGCCATACGACTCGTTCGTCGAGAACGGTGAAGATGGTGAGCTAATCGAGTACATGAGGATTAAGTCCGGCGTGGCCGATGTTCACGGAGTGGACGGCGTGCAGGGTGAGGGTGTCATCGACAGCAAATTGCTGGAAGAGTTCAAGGACAACGTGAGATACACCTTGGAAAATGACCCTGAGGAAGGAGCCGATGAGGCCACTCTGCAACGCAGGAGGCAGTTGGAACAGATCATTACGGGAGACAACGCTGAGGAGGAGTTGGAAAGGTACATCCGTGCTATGGTCAGAGAGCGCTCCACCGTGGTCAAAGACAGGGGCAAAGAGCTCCTAGGTCTATATATATATCTATATACATATTTATATATATTATTAGAACTTTACAATATAGTATATACCATTCATTGTTTAAGTTTCGGGTAATACTTTTTTTTTCCTTGTCATAACCCCAAAAATTTTCGATGCCTTTGATATAATTGAGAACAAGAAGAGTTTGCAGGTGACAAAAATCGATGATTATAGGTGTTGTGACGACAAAATGAACGCTAATATATGGGTGGCTGCTTCAGATGGTAATTTGGACCGAGTGGAACATATCCTCCGCGAGAGTAAAGGCGCCATGACCCCGCAATCCAAGGACATTAACGGCTACACTCCAATGCATGCTGCCGCCGCATACGGCCACCTGGATTTGCTGAAGAAAATGTGCAATGAGTACAATGGAGACAT

### Supplementary Note 8. Primers used for YCR051W amplification

YCR051W_fwd_BssHII: TCA**<u>GCGCGC</u>**TAATGTGTTGGACAACGACG

YCR051W_rev_SalI: CTA**<u>GTCGAC</u>**ATGTCTCCATTGTACTCATTGC

## REFERENCES

1. Frost, L. S., Leplae, R., Summers, A. O. & Toussaint, A. Mobile genetic elements: The agents of open source evolution. Nature Reviews Microbiology vol. 3 722–732 Preprint at 10.1038/nrmicro1235 (2005).

2. Rouches, M. V., Xu, Y., Cortes, L. B. G. & Lambert, G. A plasmid system with tunable copy number. Nature Communications 2022 13:1 13, 1–12 (2022).

3. Sota, M. & Top, E. Horizontal Gene Transfer Mediated by Plasmids. https://www.caister.com/hsp/abstracts/pla/05.html (2008).

4. Ausubel, F. M. et al. Current Protocols in Molecular Biology Kevin Struhl (eds.) Current Protocols in Molecular Biology. (2003).

5. Sikorski, R. S. & Hieter, P. A system of shuttle vectors and yeast host strains designed for efficient manipulation of DNA in Saccharomyces cerevisiae. Genetics 122, 19–27 (1989).

6. Myers, A. M., Tzagoloff, A., Kinney, D. M. & Lusty, C. J. Yeast shuttle and integrative vectors with multiple cloning sites suitable for construction of lacZ fusions. Gene 45, 299–310 (1986).

7. Hill, J. E., Myers, A. M., Koerner, T. J. & Tzagoloff, A. Yeast/E. coli shuttle vectors with multiple unique restriction sites. Yeast 2, 163–167 (1986).

8. Strathern, J. N. & Higgins, D. R. [21] Recovery of plasmids from yeast into Escherichia coli: Shuttle vectors. Methods Enzymol 194, 319–329 (1991).

9. Brunelli, J. P. & Pall, M. L. A series of yeast shuttle vectors for expression of cDNAs and other DNA sequences. Yeast 9, 1299–1308 (1993).

10. Frazer, L. A. N. & O’Keefe, R. T. A new series of yeast shuttle vectors for the recovery and identification of multiple plasmids from Saccharomyces cerevisiae. Yeast 24, 777–789 (2007).

11. Chou, C. C., Patel, M. T. & Gartenberg, M. R. A series of conditional shuttle vectors for targeted genomic integration in budding yeast. FEMS Yeast Res 15, 1–9 (2015).

12. Gnügge, R., Liphardt, T. & Rudolf, F. A shuttle vector series for precise genetic engineering of Saccharomyces cerevisiae. Yeast 33, 83–98 (2016).

13. Gnügge, R. & Rudolf, F. Saccharomyces cerevisiae Shuttle vectors. Yeast 34, 205–221 (2017).

14. Hohnholz, R. & Achstetter, T. Recombinant multicopy plasmids in yeast – interactions with the endogenous 2 μm. FEMS Yeast Res 19, (2019).

15. Yuan, J., Mo, Q. & Fan, C. New Set of Yeast Vectors for Shuttle Expression in Escherichia coli. ACS Omega 6, 7175–7180 (2021).

16. Gligorovski, V., Labagnara, M. & Rahi, S. J. Light-directed evolution of dynamic, multi-state, and computational protein functionalities. bioRxiv Preprint at 10.1101/2024.02.28.582517 (2024).

17. Sadeghi, A., Dervey, R., Gligorovski, V., Labagnara, M. & Rahi, S. J. The optimal strategy balancing risk and speed predicts DNA damage checkpoint override times. Nat Phys 18, 832–839 (2022).

18. Kreiss, P. et al. Plasmid DNA size does not affect the physicochemical properties of lipoplexes but modulates gene transfer efficiency. Nucleic Acids Res 27, 3792 (1999).

19. Ribeiro, S. et al. Plasmid DNA Size Does Affect Nonviral Gene Delivery Efficiency in Stem Cells. https://home.liebertpub.com/cell 14, 130–137 (2012).

20. Schakowski, F. et al. A Novel Minimal-Size Vector (MIDGE) Improves Transgene Expression in Colon Carcinoma Cells and Avoids Transfection of Undesired DNA. Molecular Therapy 3, 793–800 (2001).

21. Staal, J., Alci, K., De Schamphelaire, W., Vanhoucke, M. & Beyaert, R. Engineering a minimal cloning vector from a pUC18 plasmid backbone with an extended multiple cloning site. Biotechniques 66, 254–259 (2019).

22. Staal, J. & Beyaert, R. Extreme miniaturization in plasmid design: generation of the 903 bp cloning vector pICOt2. doi:10.1101/2023.11.29.569326.

23. Li, C. et al. Construction and functional verification of size-reduced plasmids based on TMP resistance gene dfrB10. Microbiol Spectr 11, (2023).

24. Manigat, F. O. et al. pUdOs: concise plasmids for bacterial and mammalian cells. bioRxiv 2023.07.05.547852 (2023) doi:10.1101/2023.07.05.547852.

25. Christianson, T. W., Sikorski, R. S., Dante, M., Shero, J. H. & Hieter, P. Multifunctional yeast high-copy-number shuttle vectors. Gene 110, 119–122 (1992).

26. Brachmann, C. B. et al. Designer deletion strains derived from Saccharomyces cerevisiae S288C: A useful set of strains and plasmids for PCR-mediated gene disruption and other applications. Yeast 14, 115–132 (1998).

27. Chee, M. K. & Haase, S. B. New and redesigned pRS plasmid shuttle vectors for genetic manipulation of saccharomyces cerevisiae. G3: Genes, Genomes, Genetics 2, 515–526 (2012).

28. Struhl, K. & Davis, R. W. Production of a functional eukaryotic enzyme in Escherichia coli: cloning and expression of the yeast structural gene for imidazole-glycerolphosphate dehydratase (his3). Proceedings of the National Academy of Sciences 74, 5255–5259 (1977).

29. Miozzari, G., Niederberger, P. & Huetter, R. Tryptophan biosynthesis in Saccharomyces cerevisiae: control of the flux through the pathway. J Bacteriol 134, 48–59 (1978).

30. Toh, E. A., Guerry-Kopecko, P. & Wickner, R. B. A stable plasmid carrying the yeast Leu2 gene and containing only yeast deoxyribonucleic acid. J Bacteriol 141, 413–416 (1980).

31. Lacroute, F. Regulation of Pyrimidine Biosynthesis in Saccharomyces cerevisiae. J Bacteriol 95, 824–832 (1968).

32. natMX6 Sequence and Map. https://www.snapgene.com/plasmids/yeast_plasmids/natMX6.

33. kanMX Sequence and Map. https://www.snapgene.com/plasmids/yeast_plasmids/kanMX.

34. Lin-Chao, S. Chen, W. -T & Wong, T. -T. High copy number of the pUC plasmid results from a Rom/Rop-suppressible point mutation in RNA II. Mol Microbiol 6, 3385–3393 (1992).

35. Gietz, R. D. & Akio, S. New yeast-Escherichia coli shuttle vectors constructed with in vitro mutagenized yeast genes lacking six-base pair restriction sites. Gene 74, 527–534 (1988).

36. Azoitei, M. L. et al. Computation-guided backbone grafting of a discontinuous motif onto a protein scaffold. Science (1979) 334, 373–376 (2011).

37. Reynolds, K. A., McLaughlin, R. N. & Ranganathan, R. Hot Spots for Allosteric Regulation on Protein Surfaces. Cell 147, 1564–1575 (2011).

38. Sugiura, S., Ohkubo, S. & Yamaguchi, K. Minimal essential origin of plasmid pSC101 replication: requirement of a region downstream of iterons. J Bacteriol 175, 5993–6001 (1993).

39. Mahapatra, N. R., Ghosh, S., Deb, C. & Banerjee, P. C. Resistance to cadmium and zinc in Acidiphilium symbioticum KM2 is plasmid mediated. Curr Microbiol 45, 180–186 (2002).

40. Baker-Austin, C., Dopson, M., Wexler, M., Sawers, R. G. & Bond, P. L. Molecular insight into extreme copper resistance in the extremophilic archaeon ‘Ferroplasma acidarmanus’ Fer1. Microbiology (Reading) 151, 2637–2646 (2005).

41. Gietz, R. D. & Schiestl, R. H. High-efficiency yeast transformation using the LiAc/SS carrier DNA/PEG method. Nature Protocols 2007 2:1 2, 31–34 (2007).

42. Diss, G. & Landry, C. R. Synthetic complete (SC) medium. Cold Spring Harb Protoc 2016, pdb.prot090035 (2016).

